# Integration of multi-omics data and deep phenotyping of potato enables novel insights into single- and combined abiotic stress responses

**DOI:** 10.1101/2024.07.18.604140

**Authors:** Maja Zagorščak, Lamis Abdelhakim, Natalia Yaneth Rodriguez-Granados, Jitka Široká, Arindam Ghatak, Carissa Bleker, Andrej Blejec, Jan Zrimec, Ondřej Novák, Aleš Pěnčík, Špela Baebler, Lucia Perez Borroto, Christian Schuy, Anže Županič, Leila Afjehi-Sadat, Bernhard Wurzinger, Wolfram Weckwerth, Maruša Pompe Novak, Marc R. Knight, Miroslav Strnad, Christian Bachem, Palak Chaturvedi, Sophia Sonnewald, Rashmi Sasidharan, Klára Panzarová, Kristina Gruden, Markus Teige

## Abstract

Potato is highly water and space efficient but susceptible to abiotic stresses such as heat, drought, or flooding, which are severely exacerbated by climate change. Understanding of crop acclimation to abiotic stress, however, remains limited. Here, we present a comprehensive molecular and physiological high-throughput profiling of potato (*Solanum tuberosum*, cv. Desirée) under heat, drought and waterlogging applied as single stresses or in combinations designed to mimic realistic future scenarios. Stress-responses were monitored via daily phenotyping and multi-omics analyses of leaf samples comprising transcriptomics, proteomics, metabolomics and hormonomics at several timepoints during and after stress treatments. Additionally, critical metabolites of tuber samples were analysed at the end of the stress period. Integrative analysis of multi-omics data was performed using a bioinformatic pipeline, which was established here, based on machine learning and knowledge networks. Overall, waterlogging had the most immediate and dramatic effects on potato plants, interestingly activating ABA-responses similar to drought stress. In addition, we observed distinct stress signatures at multiple molecular levels in response to heat or drought and to a combination of both. In response to all treatments, we found a downregulation of photosynthesis at different molecular levels, an accumulation of minor amino acids and diverse stress induced hormones. Our integrative multi-omics analysis provides global insights into plant stress responses, facilitating improved breeding strategies towards climate-adapted potato varieties.

**One Sentence Summary:** Integrated multi-omics analysis of high-throughput phenotyping in potato reveals distinct molecular signatures of acclimation to single and combined abiotic stresses related to climate change.

## INTRODUCTION

Improving crop resilience to climate change is a major challenge of modern agriculture (Bailey-Serres et al., 2019; Rivero et al., 2022). High-yielding crop varieties including potato (*Solanum tuberosum*), are vulnerable to heat, drought, and flooding (Benitez-Alfonso et al., 2023; Zandalinas et al., 2023; Renziehausen et al., 2024; Sato et al., 2024). These environmental stresses affect plant growth, source-sink relationships, sugar and hormone metabolism, among other processes, which in turn, negatively impact product yield and nutritional status (Lal et al., 2022). Potato is particularly sensitive to waterlogging (Jovović et al., 2021), and flooding of the fields can ruin the entire harvest within a few days. Since global warming is increasing the occurrence of such extreme weather events, crop productivity worldwide is under considerable threat (FAO, 2023). To ensure future food security, there is an urgent need for sustainable farming practices including the development of stress tolerant varieties with consistent yields (Dahal et al., 2019; Lal et al., 2022).

There is already a good understanding of how plants react to single abiotic stresses, which have profound effects on plant metabolism and development. The primary effects of abiotic stress are generation of reactive oxygen species (ROS), destabilisation of proteins and changes in enzyme efficiencies and membrane fluidity and integrity (Zhang et al., 2022). Together, these impacts reduce plant productivity through changes in photosynthetic capacity, hormone balance, transport of assimilates from source to sink as well as transport of soil nutrients and water by the roots. In addition, species-specific vulnerabilities impact agronomic productivity, such as for instance tuber initiation and tuber growth dynamics with potato. Potato tuber formation and growth is dependent on tuberisation signals produced in source leaves, such as the mobile tuberigen StSP6A (Navarro et al., 2011) that also regulates directional transport of sucrose to the developing tuber (Abelenda et al., 2019). Heat, drought and flooding trigger strong changes in gene expression and thereby strongly interfere with the regulation of flowering and tuberisation by the photoperiodic pathway. This leads to a delay in tuberisation and anomalies in subsequent tuber development such as second-growth and/or internal defects which together severely impacts marketable yields of the tuber crop (Dahal et al., 2019; Lal et al., 2022).

On the other hand, the response of plants to combined stresses is unique and cannot be extrapolated from the response to the corresponding individual stresses (Mittler, 2006). Considering the increasing occurrences of simultaneous or sequential abiotic stresses in the field, the relative lack of knowledge on multi-stress resilience is a major shortfall that hinders the ability to develop effective strategies for crop improvement. Accordingly, the question of how combinations of different stresses impact plants has recently gained a lot of interest (Zandalinas et al., 2021). Several studies on combinatorial stress effects have been performed, especially studying the physiological and molecular responses to combined heat- and drought stress in potato (Demirel et al., 2020), wheat (Manjunath et al., 2023) and tomato (Zeng et al., 2024). In nature, heat and drought often occur together, resulting in different physiological responses as compared to individual stresses. For example, under heat stomatal conductance and transpiration are increased to reduce leaf temperature, whilst under drought, stomata are closed to avoid water loss, which leads to a strongly reduced CO_2_-assimilation (Zhang and Sonnewald, 2017). The final phenotypic output in a combined stress scenario greatly depends on synergistic and antagonistic interactions between stress-specific signalling and response pathways. These interactions can be regulated at various levels (gene expression to metabolism), and on different scales (cell to system), thus resulting in complex regulatory network perturbations. Therefore, as information gained by extrapolating from studies on individual stressors is limited, it is crucial to increase our understanding of crop responses in multi-stress situations.

To this end, high-throughput phenotyping platforms and integrative omics technologies can measure molecular mechanisms at multiple levels and in multiple processes simultaneously. This can help us obtain a comprehensive understanding of the intricate dynamics of plant-environment interactions (Yang et al., 2020; Hall et al., 2022; Zhang et al., 2022). Here, advanced data integration pipelines can aid with unbiased integration and systematic extraction of biological knowledge from large multi-omics data sets (Cembrowska-Lech et al., 2023). However, despite the increasing application of high-throughput approaches in agricultural and plant research, only a handful of studies have addressed the problem of data integration from comprehensive multi-omics data (Jamil et al., 2020). Therefore, to enable molecular insights across various system levels and disentangle the intricate physiological and molecular crosstalk in the context of non-additive effects of different stress combinations, it is imperative to develop and apply multi-omics integrative approaches that leverage statistics, machine learning, and graph theory.

In this study, we aimed to increase knowledge on multiple abiotic stress responses of potato plants and to integrate this into a complex knowledge network. Therefore, a comprehensive assessment of potato responses to single and combined heat, drought, and waterlogging stress was performed. Using the cv. Désirée, a widely used moderately stress-resistant potato cultivar, we monitored dynamic changes in morphological, physiological as well as biochemical and molecular responses under stress conditions. With the application of high throughput phenotyping, multi-omics technologies, prior-knowledge and multi-level integration approaches, we identified important molecular signatures, unique to single and different stress combinations. These results can guide the development of diagnostic markers for rapid detection of stress, allowing for earlier agricultural interventions to enhance plant resilience towards abiotic stress and development of novel marker-assisted breeding programs for climate-resilient crops (Weckwerth et al., 2020; Mishra et al., 2024).

## RESULTS

### High throughput physiological phenotyping of potato in multi-stress conditions

This study aims to increase understanding of acclimation of potato plants to abiotic stresses by focusing on most likely weather (stress) scenarios in future agricultural farming: individual heat, drought and waterlogging stress, and realistic combinations thereof. We studied the cv. Desirée, a widely used moderately stress-resistant potato cultivar, as a model. Daily phenotyping using image-based sensors and regular sampling of leaf material for downstream molecular and biochemical analysis allowed better insights into the mechanisms of the stress response. Single and combined stresses were applied to plants during the tuber initiation stage at day 32 and 39 respectively, according to the scheme in Figure 1A (for more details see Supp. Table 1). Daily phenotyping of plants (control and stress conditions) was performed using the PlantScreen^TM^ phenotyping platform to assess quantitative morphological and physiological traits (Supp. Table 2), employing multiple imaging sensors (Figure 1B). Plants were light-adapted before chlorophyll fluorescence measurements to ensure steady conditions prior to the measurements.

**Figure 1.**
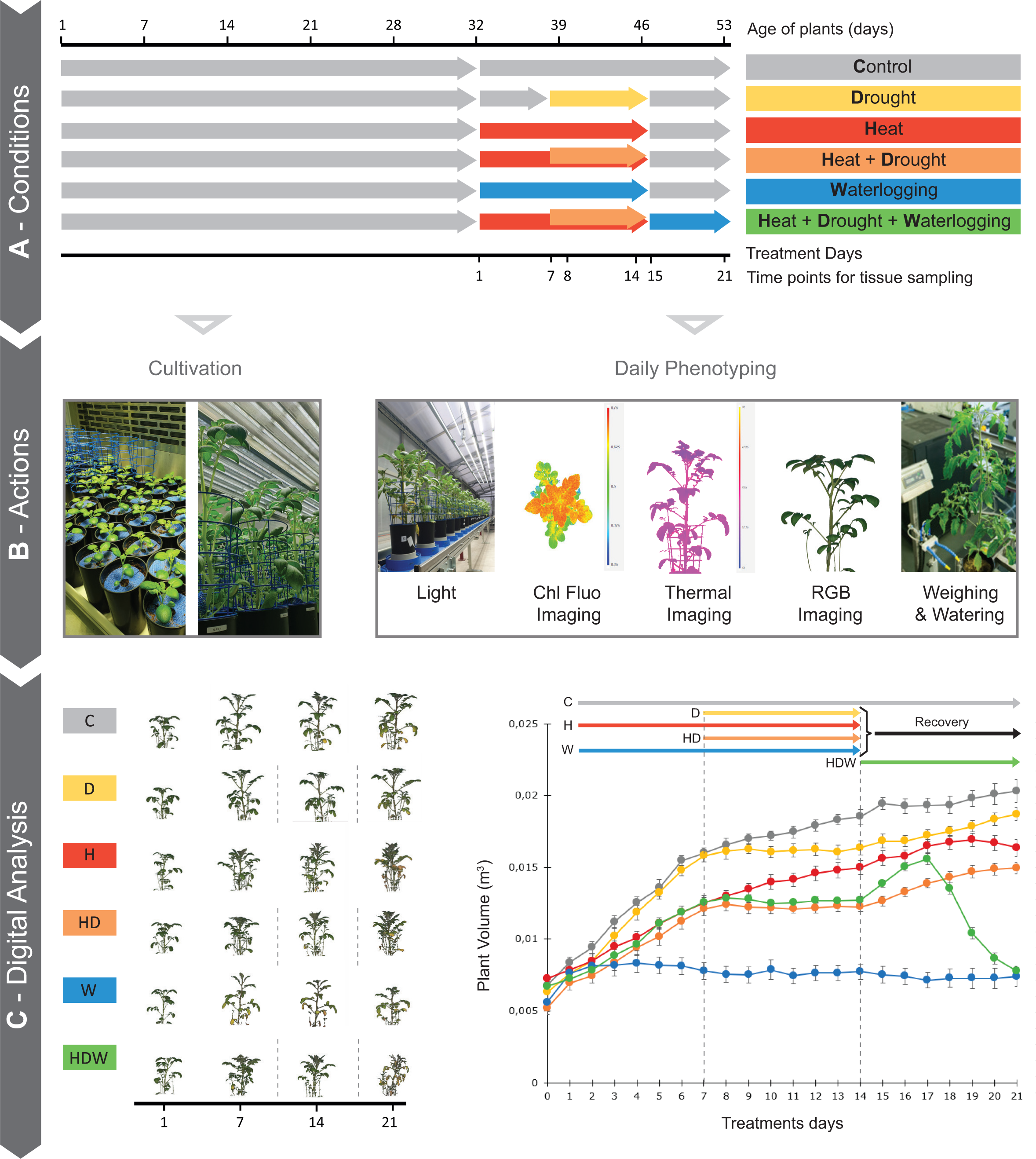
Overview of the experimental design for single- and combined stress treatments and multi-omics sampling. A) Summary of cultivation conditions. Timeline of the experimental set-up and applied stress treatments, including the recovery phase in potato cv. Desirée. Timing and duration of stress treatment and days for tissue sampling are shown. (B) Actions comprised cultivation in the growing chambers and daily phenotyping with a set of sensors using the PlantScreen^TM^ phenotyping platform at PSI Research Center. (C) Automated image analysis pipeline was used to extract quantitative traits for morphological, physiological, and biochemical performance characterization of the plants during the stress treatment and recovery phase. Side view colour segmented RGB images of plants at selected time points of tissue sampling (left panel) and daily plant volume (m^3^) calculated from top and multiple angle side view RGB images (right panel). N=6.

### Effects of single and combined stresses on potato growth and morphology

By using RGB side and top view imaging we monitored changes in plant growth dynamics during the stress treatments and recovery phases, including traits such as plant volume, area, height and compactness. The exponential growth of plants grown under control conditions was negatively impacted to various extent by different stresses (Figure 1C, Supp. Figure 1). Thus, under heat stress (H, 30°C during day, 28°C during night) the rate of biomass accumulation (plant volume) was decreased after three days similarly to drought stress (D, 30% of field capacity) (Figure 1C, Supp. Figure 1A-C). Additional morpho-physiological alterations became visible in response to heat and waterlogging (W). Heat clearly reduced plant height and caused the typical hyponastic movement of leaves (Figure 1C and Supp. Figure 1C). The negative effects of heat became more severe when combined with drought (HD, water withdrawal starting after 7 days of H) (Supp. Figure 1C). Under HD, plants phenotypically more resembled heat-stressed plants, e.g. with respect to top area and compactness, however, with a clearly more negative effect (Supp. Figure 1B, 1D). Waterlogging led to an epinastic leaf movement which was accompanied by growth arrest and significant decrease in the top area, compactness and relative growth rate (RGR) that were observed after one day (Figure 1C, Supp. Figure 1A, 1B, 1D, 1E, Supp. Table 3). In the 3^rd^ week, when the single and HD stress treatments were finished (at treatment day 15), plants recovered well from H and HD, which was clearly reflected by resumption of growth, but not from W stress (Figure 1C, Supp. Figure 1B).

Plant performance was the worst in the triple-stress condition (HDW), where 7 days of H were followed by 7 days of combined HD and 7 days of W. Interestingly, during the first three days of W that followed the period of heat and drought plants were growing very fast, but with a prolonged stress exposure, plants collapsed, as indicated from both RGB side and top view images, from the plant volume and growth dynamics as shown in RGR and other measured morphological traits (Figure 1C, Supp. Figure 1, Supp. Table 3).

### Photosynthesis is most affected by heat- and drought stress

To evaluate photosynthetic performance under single and multiple stresses, a broad range of physiological traits were extracted from chlorophyll fluorescence images and analysed (Figure 2). Top view images of the operating efficiency of photosystem II in light steady state (QY_Lss) clearly showed the negative impact of stress on photosynthetic capacity in all stress treatments, indicated by the reduction of QY_Lss, with D stress causing only a weak negative effect (Figure 2A, 2B). Moreover, steady-state fluorescence of maximum efficiency of PSII in the light (F_v_/F_m__Lss) showed a significant decrease after 3 days in W alone and when W followed a period of HD. There was also a progressive decrease in steady-state estimation of the fraction of open reaction centres in PSII in the light (qL_Lss) under whenever W was applied (Figure 2C, 2D, Supp. Table 3). Both parameters decreased even further after stress recovery following triple stress, indicating a high stress level.

**Figure 2.**
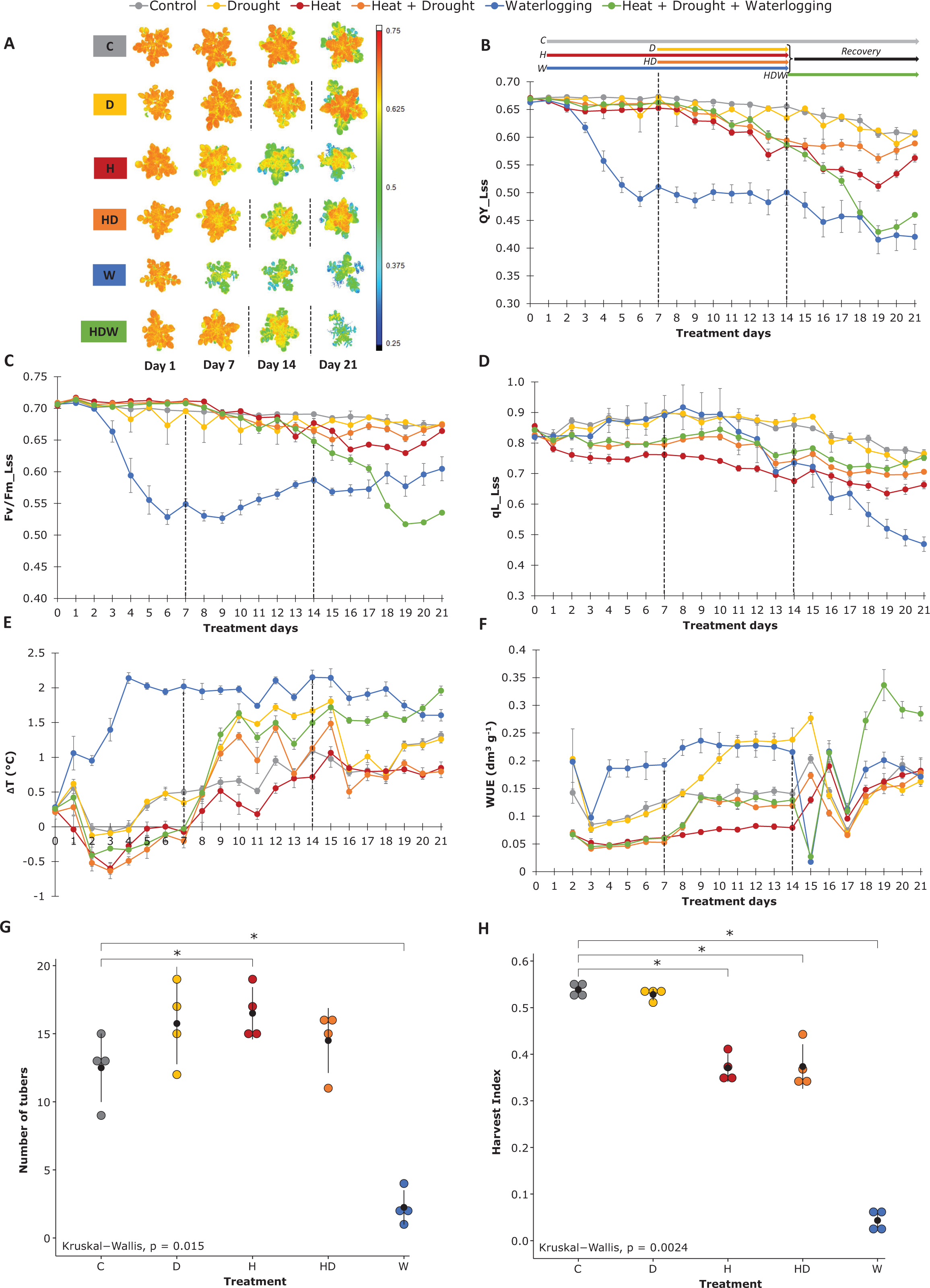
Physiological profiling using high-throughput phenotyping platforms reveals distinct responses to single and combined stresses. A) Pixel-by-pixel false colour images of operating efficiency of photosystem II in light steady state (QY_Lss) captured by kinetic chlorophyll fluorescence measurement. Images for selected time points of tissue sampling are shown. Colour coding of the treatments apply for the entire figure. Vertical dashed lines indicate the onset and end of drought. B) QY_Lss values extracted from images for each individual time point. C) Steady-state fluorescence of maximum efficiency of PSII photochemistry in the light trait based on chlorophyll fluorescence top view (Fv/Fm_Lss). D) steady-state estimation of the fraction of open reaction centres in PSII trait in light based on chlorophyll fluorescence top view (qL_Lss). E) Difference between canopy average temperature extracted from thermal IR images and air temperature measured in the thermal IR imaging unit (!::T). F) Water use efficiency (WUE) based on plant volume and water consumption. A-F) Black dotted lines reflect the initiation and removal of drought stress, respectively. See Supplementary Table 3 for Statistical evaluation of differences between groups using Wilcoxon test. G) Tuber numbers counted per plant on the last day of the experiment (Day 28 = 60 days of cultivation). H) Harvest index calculated from the total biomass and tuber weight on the last day of the experiment. G-H) Measurements, mean and standard deviation are shown (n = 4). Statistical evaluation of differences between groups is given by the non-parametric Kruskal–Wallis test (*one-way ANOVA on ranks*); p-value above x-axis, where asterisk denotes p-value < 0.05. See Figure 1A for scheme on stress treatments.

A decrease in qL_Lss was also observed after one day of H and remained constantly lower than in other conditions, while D caused no significant effect on these parameters (Figure 2D, Supp. Table 3). By applying drought in addition to heat stress, an increase in qL_Lss as compared to H alone was observed. After H and HD stress, qL_Lss values did not reach control levels till the end of the experiment at day 21 (Figure 2C and 2D), suggesting that photosynthesis is enduringly affected. In addition, changes in canopy temperature (ΔT) were deduced from the thermal imaging, while water use efficiency (WUE) was calculated based on plant volume and water consumption. The rapid increase in ΔT and WUE under W and HDW was most likely caused by prompt stomata closure (Figure 2E, 2F). The strong response remained over the entire stress period and plants did not recover from both stress treatments. A steady increase in ΔT and WUE was observed starting at three days in D, suggesting that the stress was recognised, and the plants responded by closing stomata. When D stress was released on day 15, the plants recovered immediately (Figure 2E, 2F). An opposing response was observed in H, where ΔT decreased together with an increase in water consumption, thus indicating enhanced leaf cooling (Figure 2E, Supp. Table3, 4). During combined HD stress, it appeared that the drought response became dominant over the heat responses in the physiological responses as indicated by the increase in ΔT and WUE. At the time when the drought stress was applied in addition to heat, as both parameters behaved more like drought stress responses as compared to H alone.

### Stress combinations and waterlogging have strong effects on potato yield

At the end of the phenotyping, plants were harvested to assess the total biomass accumulation and tuber yield (Figure 2G and 2H). Single H stress led to a slightly higher tuber number (Figure 2G). Harvest index was significantly reduced by H as well as by HD, while D alone did not affect final tuber yield (Figure 2H). W stress strongly inhibited tuber formation and growth and only a few tubers were formed, leading to a significant reduction in the harvest index compared to the control treatment (Figure 2H). A combination of all stress factors abolished tuber formation, reflecting the (near) lethal effect of HDW (Figure 2G, 2H).

Negative effects of the stress treatments on tubers were also observed at the metabolic level (Supp. Figure 2). Thus, starch content was significantly lower under H, HD and W stress, while drought stress alone has no negative impact. The accumulation of hexoses under H and HD may hint to an increased starch degradation and / or to a reduced starch biosynthesis. W caused a strong accumulation of almost all amino acids, most likely caused by protein degradation and a hampered metabolism (Supp. Figure 2).

### Molecular responses across omics levels reveal mechanistic insights into multi-stress acclimation

In addition to the morphological and physiological measurements (68 variables, Figure 3D, Supp. Table 1, 2), leaf samples were taken for parallel multi-omics analysis. Samples from (mature) leaves two and three were pooled, homogenized and used for further analysis (Figure 3A). For each of the treatments the fast response (one day post treatment) and the status at the end of a prolonged stress duration (7 or 14 days of stress) was investigated (sampling points see Figure 1A). While the proteome analysis was untargeted (4258 identified proteins, Supp. Table5, 6), other omics analyses were targeted comprising 14 pre-selected transcriptional marker genes involved in stress response and tuberisation, 13 phytohormones encompassing abscisic acid, ABA; jasmonic acid, JA; salicylic acid, SA; indole-3-acetic acid IAA, and their derivatives as well as 22 metabolites encompassing amino acids and sugars (Supp. Table4). To identify processes regulated on proteomics level we performed gene set enrichment analysis (GSEA, Supp. Table6).

**Figure 3:**
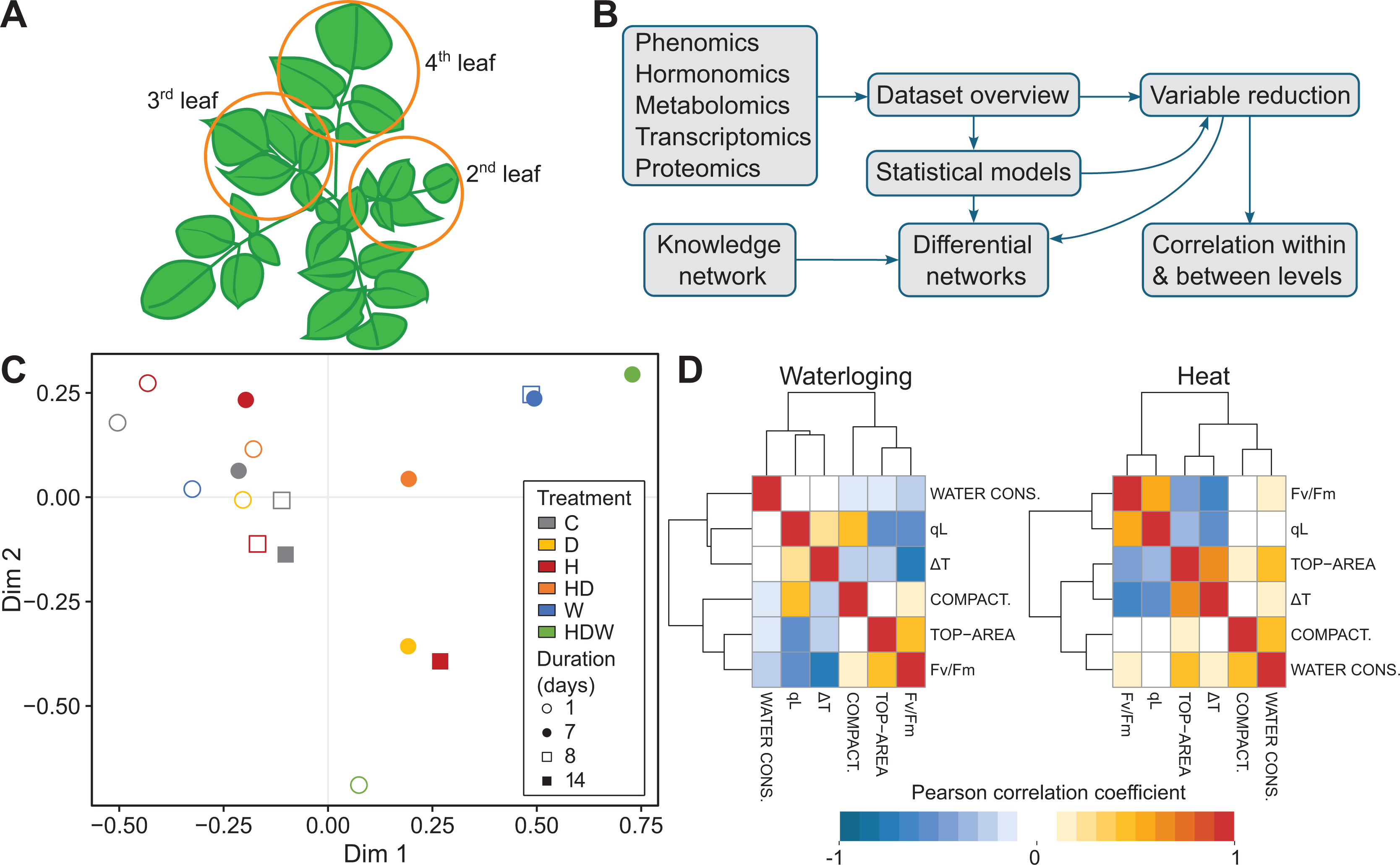
Integrated analysis of measured and generated data permits global visualization and multi-level amalgamation of potato stress responses. A) Schematics of tissue sampling protocol. 2^nd^ and 3^rd^ leaves were harvested for destructive “omics” analysis, 4^th^ leaf was used for relative water content calculation. Remaining plant tissue was quantified to obtain total above-ground biomass and tuber yield. B) Overview of data analysis pipeline. C) Dataset overview: multidimensional scaling shows combined HOW stressed plants as extremes, the centroid of each plant group is shown. D) Most informative variables from the phenomics level. Pearson correlation coefficients between them are presented as hierarchically clustered heat map in waterlogging and heat stress.

A multi-level data integration protocol was developed to investigate plant signalling/responses across the different omics levels (Figure 3B). First, we investigated data distribution by multidimensional scaling. (Figure 3C). This graph shows a clear clustering aside of samples taken after 7 and 8 days of waterlogging. Therefore, only data from the first week of waterlogging were included in further analyses, taking also into consideration that after two weeks of waterlogging all plants were severely damaged. The overview of data distribution also revealed that the most distant physiological state was that of plants exposed to triple stress (HDW): first one week of heat, followed by one week of heat combined with drought, and finally one week of waterlogging (Figure 3C). Because the triple stress treatment turned out to be very harsh and plants were severely affected in both above ground and below ground growth, we also excluded data from these samples from all further analyses. Next, we reduced the number of variables obtained on phenomics and proteomics levels to equalise numbers of variables across different analysed levels. In order to identify the most informative variables, feature selection using random forest with recursive feature elimination was conducted on the phenomics data, keeping 6 variables for downstream analysis (Figure 3D: qL, Fv/Fm, top area, delta T, compactness and water consumption). The proteomics dataset was reduced to keep only proteins that were identified as differentially abundant in any comparison of stress vs. control (135 proteins) and were functionally assigned to pathways that were studied also on other levels (36 proteins, related to photosynthesis, metabolism of sugars and amino acids, hormone metabolism and signalling, ROS signalling and stress pathways).

In addition, correlation analysis within each level of omics data was performed, revealing that these components are only weakly connected in control conditions, while in both heat or drought, they are highly correlated to each other (see e.g. for hormones and transcripts, Supp. Figure 3A, B). More severe stresses, such as the combined heat and drought stress and waterlogging, however, broke this link, suggesting a disorganisation of signalling responses. The canonical correlation analysis between components of different molecular levels similarly showed low connection in control samples. In stressed samples, blocks of components appeared to be strongly regulated, each specific to a particular stress (Supp. Figure 3C).

Variables measured on different omics levels were integrated into a metabolism and signalling cascade-based knowledge network to capture events at the molecular level (Figure 4A). Finally, we superimposed the measured data onto this mechanistic knowledge network and visualised them in parallel for all omics levels per each analysed condition compared to control (Figure 4B, Supp. Table 3). This provides a comprehensive overview of how these stresses rewire biochemical pathways and physiological processes. These networks were used for interpretation of processes in single and combined H and D stress as well as for waterlogging (W) and are described in the subsequent sections.

**Figure 4:**
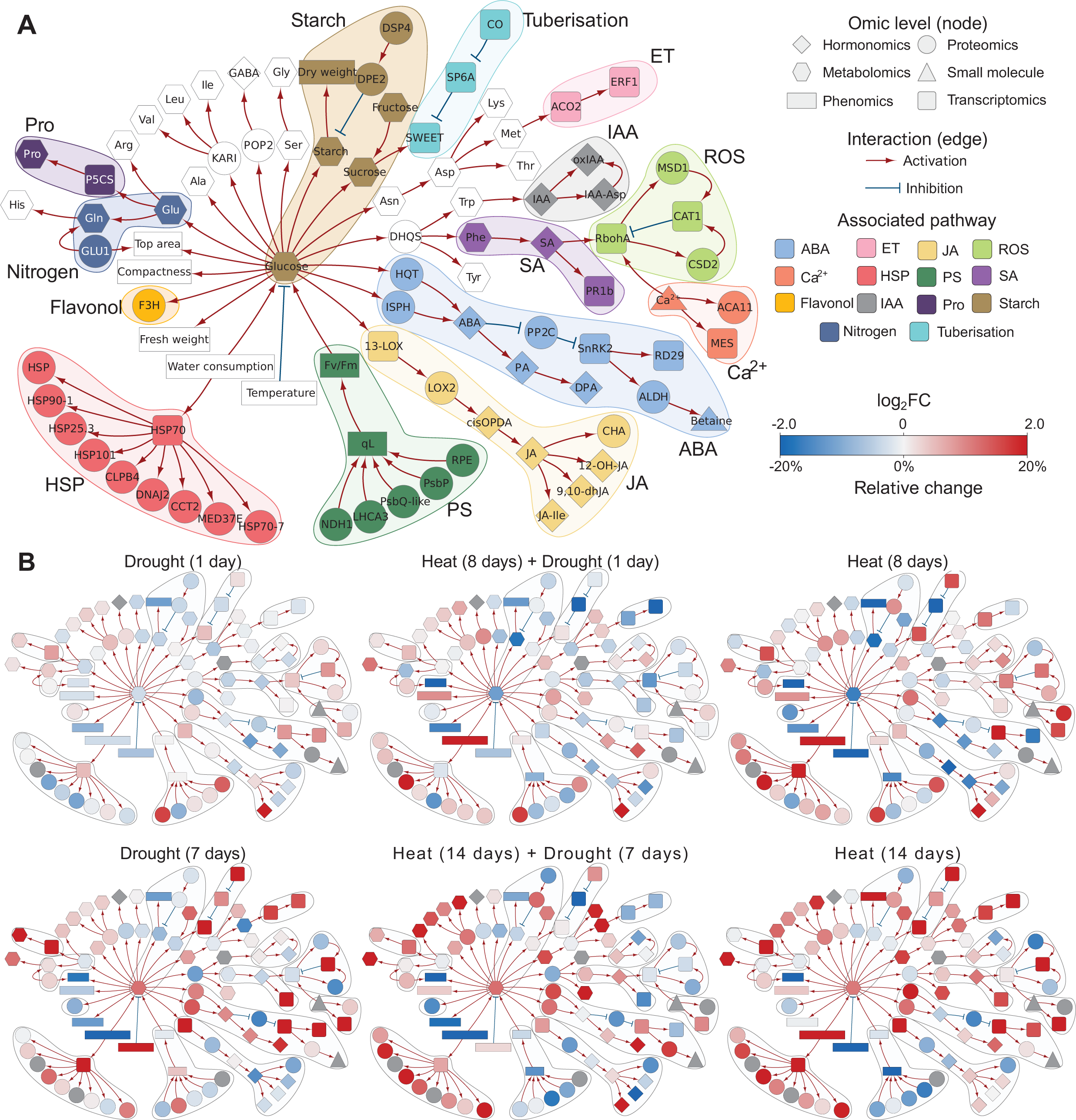
Integration of multi-omics data in a knowledge-based metabolic and signalling network. A) Structure of knowledge network. Individual studied components are coloured according to their function in different pathways. B) To compare the effects of different stresses on the overall state of the plant, we overlaid the knowledge networks with measured changes in component concentration. Nodes are coloured by log2 fold changes (red – increase in stress compared to control, blue – decrease in stress compared to control, grey – measurement not available) shown for two time points: sampling day 8 and sampling day 14 for the different stress treatments, days of stress treatment are given with each network (for more details of the set up see Figure 1A).

### Metabolic and molecular responses to individual and combined heat- and drought stress exhibit combinatorial and distinct signatures

While heat stress was effective immediately, drought stress, applied by water withdrawal on day seven in our set-up, became effective gradually within three days, visible by an increase in the ΔT values (Figure 2E). Similar to previous reports (Demirel et al., 2020; Zaki and Radwan, 2022), we found that Desirée was moderately drought tolerant and exhibited only minor morphological and physiological responses at the moderate stress level applied in our study (30% water capacity of the soil) and the plants fully recovered when the stress treatment was finished. The potato plants clearly responded to elevated temperatures (heat stress, H) with morphological adaptation like the upward movement of leaves (Figure 1A), which is part of thermo-morphogenic responses (Quint et al., 2016). Previous work showed that heat stress caused an altered biomass allocation between shoots and tubers of potato plants, with less assimilates allocated to developing tubers (Hancock et al., 2014; Hastilestari et al., 2018), leading to decreased tuber yield (Figure 2G) and starch accumulation and a decreased harvest index which was also seen in our study (Figure 2H and Supp. Figure 2).

To investigate the effect of heat and/or drought stress on leaf carbohydrate metabolism, contents of soluble sugars and starch were measured (Figure 5A). While sucrose levels did not change (Supp. Table 3, there was about twofold increase in the amounts of fructose and glucose after 14 days of H and at day 7 of D and HD. The soluble sugars may act as osmo-protectants under these stress conditions. In addition, we found that protein levels of enzymes involved in glycolysis or sucrose degradation were upregulated in H (at 7 days) and combined HD (day 14) as indicated by gene set enrichment analysis which summarizes the complex proteomics data set (Figure 5D and Supp. Table 6). This may indicate an increased demand for energy and/ or building blocks for stress defense responses. Under these conditions also less starch accumulated, visible at day one and eight in H and at day one HD, suggesting lower carbon availability due to a disturbed photosynthesis and/or an increased demand for other processes. Expression of the sugar efflux carrier *SWEET11* was upregulated at the end of both H and D stress presumably to maintain sucrose loading into the phloem and carbon allocation to sink tissue to counterbalance the decreased carbon assimilation rate of source leaves. Together with the reduced photosynthetic efficiency, that was measured, these data imply that a lower amount of carbon is assimilated, especially under H and HD, and available for allocation to sink organs, such as growing tubers. This most likely contributes to the impaired tuber growth and lower tuber starch accumulation found under these conditions (Supp. Figure 2).

**Figure 5:**
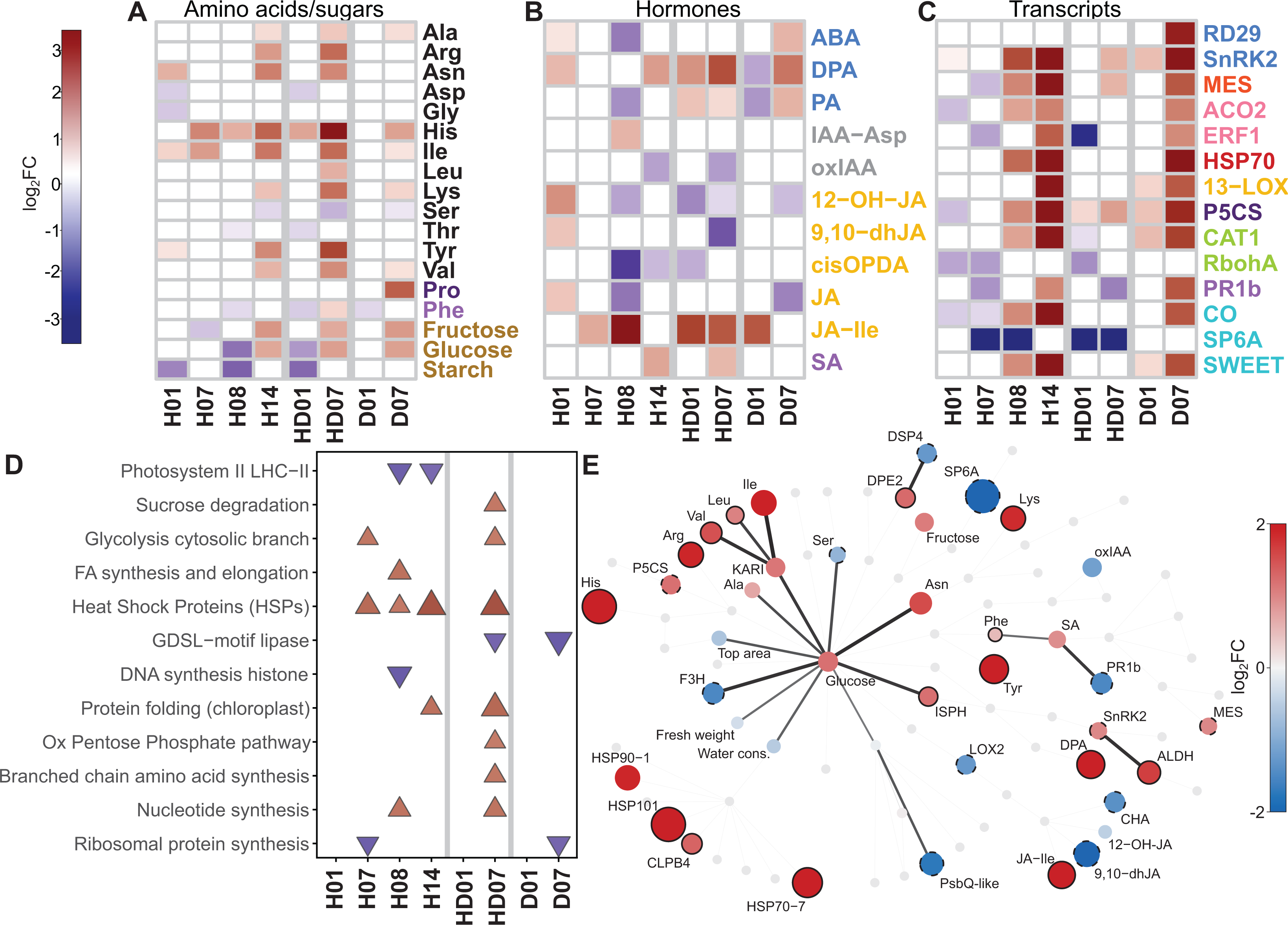
Combined heat and drought stress trigger distinct responses compared to each individual response. Additive effect of combined stress is most pronounced for branched chain amino acids accumulation and JA signalling response. A-C) Heatmaps showing log2FC (FDR p-value < 0.05) in individual stress heat (H) or drought (D) stress in comparison to combined one (HD) for targeted molecular analyses. Label colours indicate pathway associated with each molecule as in the Knowledge network (see Figure 4 for legend). A) Changes in metabolite levels. B) Changes in hormone levels, and C) Changes in selected stress-related transcripts. D) Changes observed on proteomics level. Results of Gene Set Enrichment Analysis (FDR q-value < 0.1) are shown. For more information see Supp. Table 6. C) Biochemical knowledge network showing changes under combined HD stress at day 14 (treatment day 7). In this version of knowledge network, only nodes that were significantly differentially expressed (vs. control conditions) are coloured and the connections between two differentially expressed nodes are coloured black. Node full black border indicates molecules with higher expression levels in HD compared to H and/or D alone. Dashed black border indicates molecules with lower expression levels in HD compared to H and/or D alone (difference of log2FC > 0.5).

One key player stimulating tuberisation and tuber growth is the tuberigen *SP6A*. Its expression was downregulated during the first week of H and in combined HD (Figure 5C). During longer heat exposure, expression levels of *SP6A* were similar to control levels, but remained low in HD. Drought alone had little effect on *SP6A*, which is consistent with the low impact on final tuber yield. The transcriptional regulator protein *Constans-like 1* (CO) was described to act as a negative regulator of *SP6A* expression (Abelenda et al., 2016). *CO* was upregulated within 7 days of drought stress (D), and it increased with duration of heat, but was unaffected by HD combination (Figure 5C). Hence, in our experiment, the transcript levels of *CO* did not always change in the opposite direction as *SP6A*, assuming additional regulatory mechanisms may act under stress conditions.

Considering the changes in amino acids, the most striking finding was the strongly elevated levels of histidine (His) in all three stress treatments, with the highest amounts detected in combined HD stress (Figure 5A). This was accompanied by a significant increase of many (minor) amino acids, in particular iso-leucine (Ile) and other branched chain amino acids (BCAs). This observation was in line with previous reports on combined heat- and drought stress in potato (Demirel et al., 2020), although it’s cause and importance needs further investigations. Accordingly, at the proteome level, proteins involved in BCA synthesis were significantly enriched among the ones with increased levels in stress (Figure 5D, 5E, Supp. Table 5, 6).

Proline is known as an important regulator of osmotic potential that protects cells by stabilizing proteins and scavenging of reactive oxygen species. Its amount was elevated at day 7 of D stress (sampling day 14) (Figure 5A). This was consistent with increased transcript levels of *P5CS* (*Pyrroline-5-carboxylate synthase*), the key enzyme for proline synthesis, and of *RD29* (*Responsive to Desiccation*), both being well-known (drought-) stress marker genes. Consistently, levels of ABA, the key phytohormone that induced stomal closing, proline accumulation and other drought stress responses (Cutler et al., 2010; Zhang et al., 2022), were elevated after 7 days of D but were clearly reduced in H, while no changes were detected in HD stress. Interestingly, the levels of phaseic acid (PA), and dihydrophaseic acid (DPA), two breakdown products of ABA, were lower at one day of D but significantly higher after 7 days in D. The strongest accumulation of DPA levels was detected in the HD treatment, in which DPA levels were elevated already after one day and further increased until the end of the treatment (at day 7). The elevated levels after one day of HD can be explained by the experimental setup, in which the D treatment started after 7 days of H (that also resulted in DPA accumulation). However, the strong accumulation of ABA breakdown products under D and to even higher level in HD are in line with their suggested role in long-term stress acclimation. It was shown that PA, the first degradation product of ABA and precursor of DPA, does also activate a subset of ABA receptors (Weng et al., 2016). Because ABA has a very short half-live, it was suggested that the long-lived PA could prime plants for enhanced responses to future drought (Lozano-Juste and Cutler, 2016).

The phytohormone JA is another typical stress hormone known to be involved in many biotic but also abiotic stress responses (Wasternack and Feussner, 2018). The biologically active form is jasmonyl-L-isoleucine (JA-Ile) and we found strongly increased levels of JA-Ile under H, D and HD conditions, (Figure 5B). 12-Hydroxyjasmonic acid (12-OH-JA) is a by-product of switching off JA signalling with weak signalling activity (Nakamura et al., 2011). It was also described to function as tuber-inducing factor in potato (Yoshihara et al., 1989). Under H, amounts of 12-OH-JA as well as free JA switched from higher amounts measured at day one to lower levels at day 8, and decreased further till the end of the experiment. Also, *cis*-12-oxo-phytodienoic acid (*cis*-OPDA), the biochemical precursor of JA, was detected at much lower levels on day 8 and 14 in H. Cis-OPDA was also reduced at the start of the combined HD (day one) treatment, most likely because of prior heat treatment. Altogether, this indicates a strong upregulation in the last step of conjugation for the synthesis of JA-Ile in H, D and HD.

The accumulation of ROS is a detrimental by-product of photosynthesis and other pathways under stress conditions. Accordingly, the detoxification of ROS by different enzymes such as catalase or superoxide dismutase together with induction of Ca^2+^ signals is a typical response emerging from stressed chloroplasts (Stael et al., 2015). In line with that, we measured increased transcript levels of catalase (*CAT1*) and a methyl esterase (*MES*), which was selected as Ca^2+^ signalling marker gene (Marc Knight, unpublished data) at the end of the drought and heat treatment (sampling day 14). The transcript levels of *pathogenesis-related protein 1b1* (*PR1b*), a biotic stress as well as drought and salt stress marker (Akbudak et al., 2020) were first lower in H but increased from day 8 to 14 in H and at day 7 in D (Figure 5C). A similar response was seen for the chloroplast-localized lipoxygenase *13-LOX*, which is a well-known marker gene for different stresses, especially chloroplast generated ROS (Bachmann et al., 2002) suggesting increased ROS formation and stress level with stress duration. Strikingly, the expression of these genes was less induced or even inhibited when H and D were combined, which may indicate the activation of opposing signalling pathways

Heat stress, but also other stresses, induces the production of heat-shock proteins (HSPs), which is a very conserved process in all organisms. HSPs act as molecular chaperones and play an important role in maintain the cellular homeostasis and the proteome by supporting protein folding, preventing misfolding or by assisting in the degradation of irreversibly damaged polypeptides (Sato et al., 2024). At transcript level, we observed clearly elevated levels of *HSP70* transcripts after single H and D stress. Under H this was accompanied by an accumulation of numerous HSP proteins as indicated by their significant enrichment among the identified proteins in the proteome approach (Figure 5D, 5E). This effect was similarly pronounced in combined HD stress (Figure 5E) with a strong enrichment of HSP70, 90, 101 involved in heat stress as well as in assisting folding (Figure 5D). The category “protein folding” comprises mainly HSP70 and 60 group members with many of them that are present in the chloroplast, where they are participate in the repair of photosystem (PS) II components, but also protect proteins such as RuBisCo. In fact, our physiological data indicate a disturbance in the electron-transport chain through PSII under heat. For example, we saw a strong decrease of PSII efficiency under H stress (Fig. 2B). The negative effect of H stress is also reflected in lower abundance of photosystem II proteins in the gene enrichment analysis (Figure 5D). More specifically (Supp. Table5), there were reduced amounts of the PsbQ and PsbP subunits of the oxygen-evolving complex.

Overall, we do see specific stress responses to heat and drought, but also to a combination of both. This becomes also visible in Figure 5E illustrating the signature which is caused by combined heat and drought stress in biochemical pathway view (knowledge network overlaid with multi-omics data). The responses to HD only partly overlap with single D stress, e.g. for the accumulation of DPA. However, for most measured metabolites the patterns are similar to heat, but changes to HD were more pronounced (Figure 5E). Here, we cannot exclude that this domination by heat was linked to rather the mild drought stress level. Interestingly, the transcriptional changes of selected stress-related enzymes were weakest in HD stress combination pointing to a redirection and rearrangement of signalling pathways compared to individual stress factors as suggested by other studies (Zhang and Sonnewald, 2017).

### Comprehensive insight into molecular processes mediating the extreme waterlogging sensitivity of potato

Despite being documented as a highly flood-sensitive species, an in-depth characterization of flooding-induced stress responses in potato is sparse (Jovović et al., 2021). The waterlogging sensitivity of potato was evident in the HTP data, with several morpho-physiological traits related to plant performance being negatively impacted following stress imposition (Figure 1C, Figure 2). This included: leaf epinasty, decreased biomass accumulation and shoot elongation, impaired photosynthesis and stomatal conductance as well as a dramatic reduction of tuber yield (see Figure 1,2).

Waterlogging significantly affected primary metabolic pathways as reflected in an increase in soluble sugars and free amino acids (Figure 6A). We also observed changes in the expression of stress-associated genes and hormones, thus highlighting potential mechanisms involved in waterlogging acclimation. This include the increase of ABA (ABA, PA, DPA) metabolism and response (*RDB29),* as well as the upregulation of the ethylene (ET) biosynthesis gene, *ACO2,* and the ROS-producing enzyme, *RbohA.* Waterlogging also led to accumulation of different JA metabolites (9,10-dHJA, cisOPDA, and JA-Ile) together with the upregulation of 13-LOX, an enzyme involved in JA metabolism. The strong ABA signature observed in waterlogged plants prompted us to compare waterlogging and drought responses (Figure 6B). This revealed common stress-associated responses (e.g.: the induction of *ACO2*, *RD29, HSP70*) and a much stronger ABA response in waterlogging relative to drought. Another notable observation was the upregulation of the tuberigen signal, *SP6A* after 1d of waterlogging coinciding with the downregulation of its negative regulator *CO* (Figure 6C).

**Figure 6:**
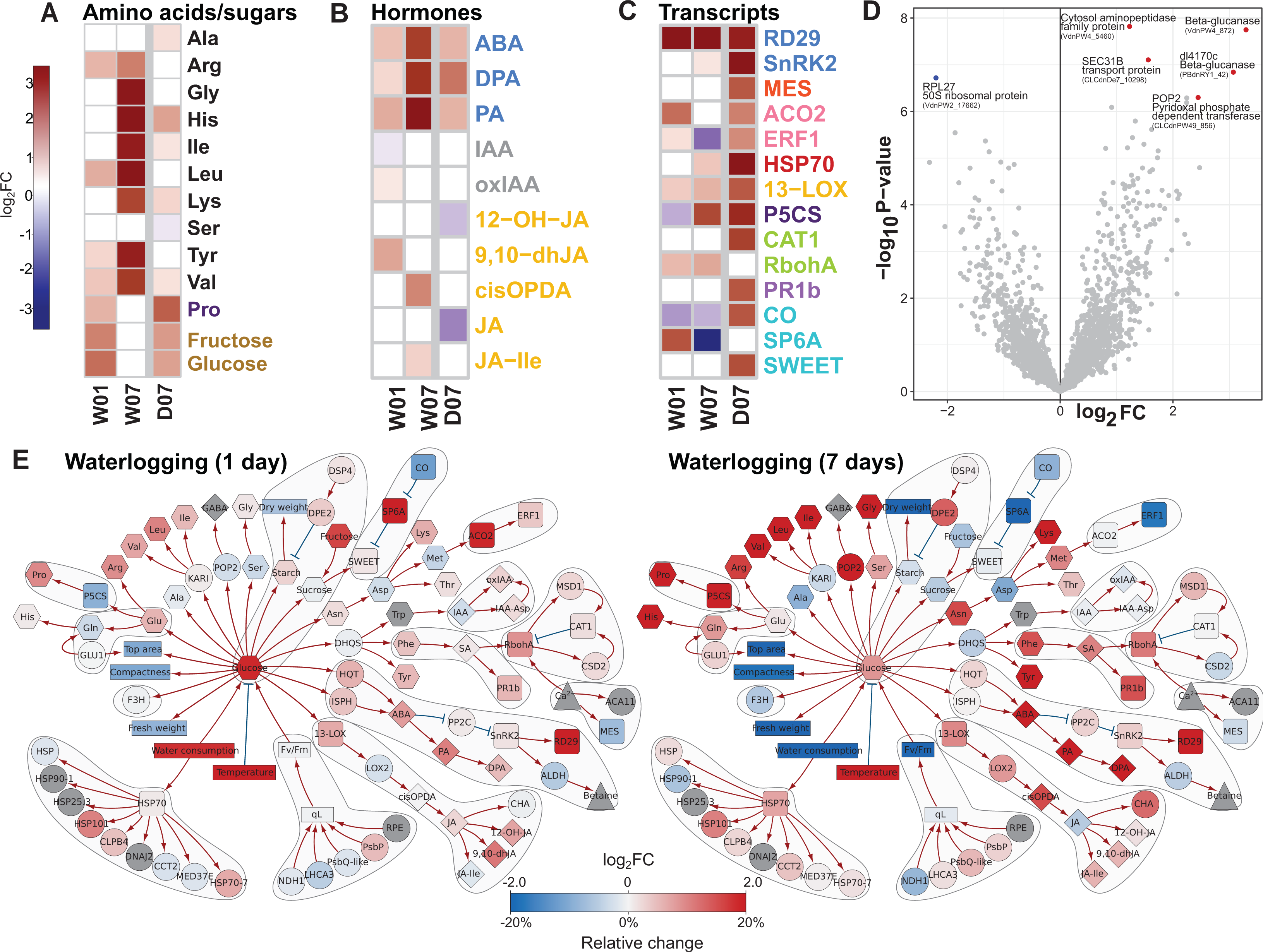
Waterlogging triggers drought-stress like molecular responses in potato. A-D) Heatmaps showing log2FC (FDR p-value < 0.05) changes in A) metabolite levels, B) phytohormones, C) selected stress-related transcripts. D) Volcano plot of differential proteomics analysis at day 7. Proteins with FDR p-value < 0.05 shown as blue (downregulated) and red (upregulated) dots. For more information see Supp. Table 5. E) Knowledge network of waterlogging stress at day 1 and day 7 (unfiltered, colour range [-2, 2]). For legend see Figure 4.

Proteomics analyses of waterlogged plants revealed mild effects. Six differentially enriched proteins were identified in response to prolonged (7 days, W07) waterlogging treatment. Among the strongly upregulated proteins, were a LAP2-like protein (VdnPW4_5460) which encodes an aminopeptidase, two glucan endo-1,3-beta-glucosidases (VdnPW4_8729, PBdnRY1_427), a POP2-like protein encoding a GABA transaminase and a SEC31B-like protein, previously described as a component of coat protein complex II (COPII), involved in vesicular transport from the endoplasmic reticulum (ER) to (Li et al., 2021). Waterlogging led to the downregulation of the chloroplast ribosomal protein RPL27-like, demonstrated to be important for protein synthesis (Figure 6D).

The multi-level integrative analyses enabled visualization of the progression of stress symptoms in waterlogged plants. In comparison to one day of waterlogging, molecular responses to prolonged waterlogging stress (7 days, W07) displayed a distinct signature (Figure 6E). While ABA-, JA- and ROS-biosynthesis and accumulation of free amino acids was further increased, we observed that prolonged waterlogging led to a general inhibition of the tuberisation process (i.e. *SP6A* downregulation). In addition, genes related to ethylene biosynthesis and response were not regulated (*ACO2)* or downregulated (*ERF1),* thus suggesting temporal control of ethylene signalling. Despite representing opposite ends of the water stress spectrum, waterlogging and drought, elicited significantly overlapping responses, notably related to ABA metabolism and proline accumulation (Figure 6A, 6B, 6E).

## DISCUSSION

Despite its outstanding importance as a major food crop, research into the vulnerability of potato to abiotic stresses lags that of other staple crops. In recent years, potato yields have been significantly affected by heat, drought, and flooding, often occurring sequentially or simultaneously (Dahal et al., 2019; Jovović et al., 2021; von Gehren et al., 2023). Considering the increasing occurrence of these extreme weather events, this knowledge gap needs to be urgently addressed. In this study, we leveraged the power of several omics techniques and their integrated analysis to build a comprehensive global picture of potato responses to single and combined heat, drought, and waterlogging stress.

### Leveraging multi-omics data integration to capture the complexity of biological systems

Several tools have already been developed for integrative analysis of multi-omics data (Joshi et al., 2024). Most broadly used are the mixOmics package (Rohart et al., 2017; Singh et al., 2019), integrating datasets based on correlations, and the pathway visualisation tool PaintOmics (Liu et al., 2022). In this study, however, we integrated five omics-level datasets. Such complex datasets have rarely been analysed, even in medical research (Lee et al., 2019), as most studies combine only two to three omics datasets (Ployet et al., 2019; Lozano-Elena et al., 2022; Núñez-Lillo et al., 2023; Núñez-Lillo et al., 2024; Sinha et al., 2024). Since existing tools were not directly suitable for our needs, we developed a novel pipeline harnessing the potential of both integrative and visualization approaches. An additional step, based on machine learning. was introduced to reduce the number of variables, in particular of phenotypic physiological data. This reduction of variables was especially important for the correlation analyses across omics levels, where we kept only the most informative variable. In the first step we performed statistical modelling and correlation analyses, which provided partial overview of events. In the second step, mechanistic insights were obtained by generating a customised biochemical knowledge network. Our network was constructed based on knowledge extracted from different databases, most notably the Stress Knowledge Map (Bleker et al., 2024) and KEGG (Kanehisa et al., 2017), as well as from literature, to integrate all components that were kept after variable selection. The obtained biochemical knowledge network enabled a comprehensive overview of events at the pathway level and led to the identification of mechanistic differences occurring in response to different stresses. The developed pipeline is thus highly useful for integration and interpretation of complex datasets in future studies and can also be applied to other species.

### Integrative omics provides global insights into potato abiotic stress responses

While multi-omics approaches have been successfully applied in numerous crop species to better understand abiotic stress responses, our study is the first to do so in potato. We subjected the cultivar Désirée to waterlogging, drought, heat, a combination of heat and drought, and triple stress combination encompassing all three. Across each stress treatment, detailed morpho-physiological traits were measured, with a subset of plants sampled for the probing of a diverse array of molecular stress markers, hormones, metabolites, and proteome analyses across several time points.

In general, stress combinations appeared to be more detrimental to the plant performance than individual stress applications. The combination of H, D and W led to a rapid decline in plant viability and eventually most plants died.□ However, all individual stress factors caused a reduced plant growth, had a negative impact on photosynthetic assimilate production, and both heat and waterlogging stress impaired tuber yield (Figure 7) and tuber starch accumulation (Supp. Figure 2). Considering all stress responses, it turned out that the cultivar Desirée was less affected by the applied drought stress indicating that it is quite resilient to drought as suggested previously (Demirel et al., 2020). A combination of heat and drought caused stronger growth retardation than both stresses alone with drought responses overwriting heat adaptations.

**Figure 7.**
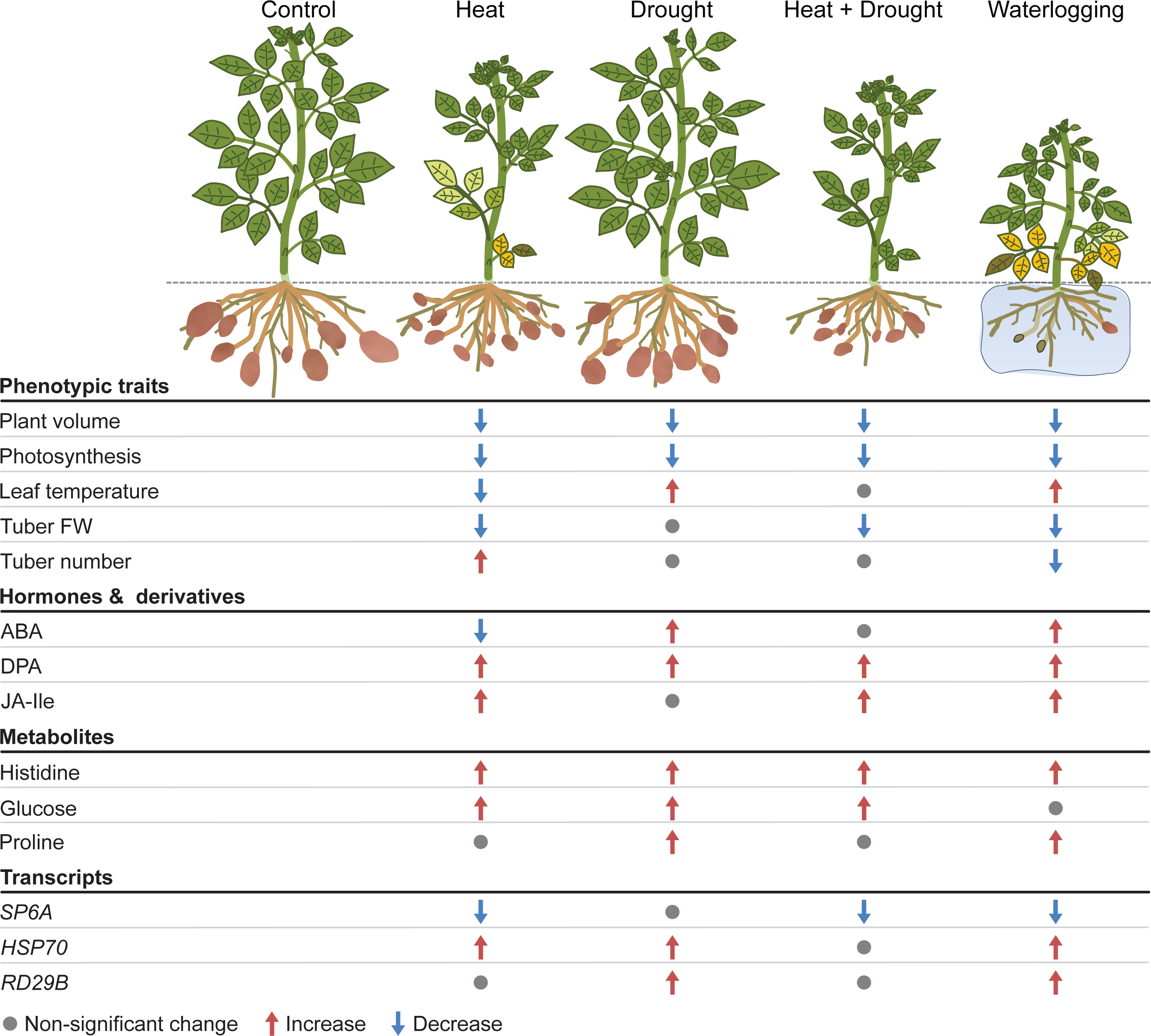
Schematic summary of multilevel responses to single and combined heat, drought and waterlogging stresses. Selected variables from each level are shown. Summary of molecular responses (hormones, metabolites and transcripts) was based on the comparisons illustrated in Figures 5 and 6. Summary of morphophysiological responses were based on the data from the last day of the experiment (Day 28), which includes tuber information. Proteomics data set is not included here due to the small dataset of differentially expressed proteins in the waterlogging treatment. Degree of increase or decrease is not specified.

Nevertheless, despite the apparent mild drought phenotypes, a clear drought-associated signature (accumulation of ABA, sugars, proline, histidine and most stress-induced transcripts) confirmed the activation of stress responsive pathways, particularly at day 7, which contributed to stress acclimation (Figure 5). Thus, there was a clear activation of the ABA response pathway, as seen by an increase in the content of the hormone and its degradation products, as well as proline, and the increased expression of ABA-responsive marker genes *SnRK*, *P5CS*, *RD29*, leading to the corresponding physiological responses, e.g. a decreased water-use and leaf temperature caused by stomata closure. Particularly interesting was the accumulation of the ABA catabolite DPA after longer drought stress. The precursor of DPA, PA was suggested to have an important role in priming for increased resilience to future drought stress in Arabidopsis (Lozano-Juste and Cutler, 2016). It is believed that DPA does not trigger ABA responses, but that has to our knowledge not been studied in potato. Hence, it might be that DPA acts as priming signal for stress acclimation and resilience in potato.

Compared to drought, heat stress had a stronger impact on Desirée plants at all levels from growth to photosynthesis and yield. Thermomorphogenesis is a well-described morphological response to elevated temperature stress comprising shoot elongation and hyponastic movement of leaves which together with an increased transpiration are seen as an acclimation to increase ventilation and to cool the aboveground part (Quint et al. 2016). In our study we clearly observed the hyponastic movement of leaves and stomatal opening, indicated by a decrease in ΔT (Figure 7). This physiological response was accompanied by a decreased amount of ABA. In contrast, the heat-mediated shoot elongation that has been seen in other potato varieties was not visible in Desirée (Hastilestari et al., 2018; Tang et al., 2018). Instead, the plant height of Desiree plants was reduced under elevated temperature suggesting cultivar-specific difference that could be exploited in further studies to entangle different morphological stress adaption mechanisms in potato. In *A. thaliana*, the thermomorphogenetic hypocotyl elongation is tightly linked with an increase in auxin levels and is mediated by the transcription factor Phytochrome interacting factor 4 (PIF4) (Quint et al. 2016). Consistent with the morphological response, we did not find significantly altered levels of the phytohormone IAA in leaves of Desiree plants in response to heat (Figure 5B).

High temperature treatment negatively affects photosynthetic capacity, particularly the efficiency of photosystem II, which is in line with results of other studies (see (Mathur et al., 2014) for review), and this was confirmed here at both physiological and proteomic level. In a previous study by (Hancock et al., 2014), which also used cv. Desiree, the net CO_2_ assimilation was even higher under elevated temperatures than in control conditions. This difference might be related to the different set-up, as in the latter study the night temperature was kept at 20°C, while here it was adjusted to 28°C, suggesting that a low night temperature is important to maintain photosynthetic activity. The heat-induced impact on photosynthetic capacity most likely caused a lower production of assimilates, indicated by the decreased amount of transitory starch in leaves. Concomitantly, contents of hexoses were found to be increased consistent with earlier studies (Hastilestari et al. 2018). This increase may contribute to the osmo-protection of cells and provides energy for the costly heat stress response such as the formation of heat shock proteins (Guihur et al., 2022). A massive accumulation of heat shock proteins was found after one week of heat stress, together with elevated levels of *HSP70* transcript levels at day 8 (Figure 5C). Although energy-demanding, the induction of HSP is important for the cellular homeostasis and maintenance of growth and metabolism at elevated temperature. This is well demonstrated by transgenic potato plants with increased expression of a beneficial allele of HSC70, that exhibit improved heat stress tolerance (Trapero-Mozos et al., 2018).

A downregulation of photosynthesis is a typical stress response to prevent potential damage, for example caused by ROS. This has strong implications on plant growth and yield and is therefore regulated at various levels including light-harvesting and electron transport with high implications for crop improvement (Kromdijk et al., 2016), particularly under stress conditions (Grieco et al., 2020). The resulting lower photosynthetic capacity together with an increased energy demand for stress defence reduces the amounts of assimilate that can be translocated toward the developing tubers and its availability for storing starch. Beside assimilates, molecular signal stimulating tuber development and growth. One important regulator is *SP6A* which was downregulated at transcript level in our studies, similar to previous ones (Hancock et al., 2014; Lehretz et al., 2020; Park et al., 2022; Koch et al., 2024). Stem-specific overexpression can overcome heat-mediated yield reduction and enhances delivery of assimilates towards tubers.

Looking at plant hormones, we observed changes of stress hormones like SA and JA, that traditionally have been associated with biotic stress responses. Quite striking in this context was the increase in amount of JA-Ile under heat, drought and the combination of both (Figure 5B). This is consistent with previous observations reporting that JA has a positive effect on thermotolerance in Arabidopsis (Clarke et al., 2009; Balfagon et al., 2019). Heat stress increased levels of OPDA, JA and JA-Ile and application of 5µM methyl-jasmonate improved cell viability (Clarke et al., 2009). Using mutants this study also showed, that JA acts in concert with SA in conferring thermotolerance. Moreover, an accumulation of JA-Ile was also observed in Arabidopsis under drought (Yoshida and Fernie, 2024), and increased levels of JA-Ile by overexpression of *JAR1* resulted in improved drought stress tolerance, however resulted in stunted growth (Mahmud et al., 2022). Detailed analyses how levels of JA and its derivates as well as biosynthesis and signalling components changes in response to stress are still missing in potato. Therefore, but a deeper understanding of regulatory factors is required, particularly the crosstalk with other hormones and the impact on plant growth. However, a tight modulation of JA metabolism seems as promising target for future engineering of abiotic stress tolerance in potato (Bittner et al., 2022).

### Extreme sensitivity to waterlogging in potato – integrative -omics highlights commonalities with drought

Our study provides the first detailed insight into the molecular responses underlying the high vulnerability of potato to waterlogging. Water saturation imposes rapid oxygen deficiency in the soil, thus impairing root respiration and function. Plant survival in flooded soils involves various morphological and metabolic responses to either escape or cope with hypoxia, which involve acclimation responses in roots but also in aerial organs (Sauter, 2013; Leeggangers et al., 2023).

The data showed that plant growth and performance were more drastically affected by waterlogging as compared to H, D and HD treatments. In addition, HTP data suggests that waterlogging had a dominant effect even when applied after a previous combined exposure to heat and drought (HDW) (Figure 1C and 2). When applied as a single stress, detrimental effects on plant performance increased over time. Waterlogging dramatically impairs root conductance and water and nutrient uptake, causing tissue dehydration and wilting. This triggers water-saving responses such as stomatal closure and epinasty, which were reflected in increased leaf temperatures and reduced plant compactness, respectively (Figure 1, 2E, suppl. Figure 1). Epinastic leaf movement, a common waterlogging response in Solanaceae, is thought to reduce photosystem damage by irradiation and transpiration (Jackson and Campbell, 1976; Geldhof et al., 2023). Both stomatal conductance and epinasty are regulated by the pivotal flooding signal ethylene (Leeggangers et al., 2023). While ethylene levels were not measured here, the analyses of synthesis genes (i.e.: *ACO2)* suggested the activation of ethylene production in waterlogged shoots. Ethylene is also known to trigger *RBOH* expression and can act synergistically with ABA to reduce stomatal conductance (Zhao et al., 2021). ABA is also considered to signal root stress during waterlogging (Jackson and Hall, 1987; Zhao et al., 2021). We observed both the activation of ABA signalling and ABA accumulation and together with increased levels of proline, this is consistent with a strong drought signature (Figure 6A, 6B, 6E). While paradoxical, waterlogging is known to elicit shoot drought responses. As root function in hypoxic soil ceases, it triggers a ‘drought like’ response in the shoot with the similar goal to trigger water saving measures. A focus on this ABA and drought-mediated regulatory network might thus be an attractive target for probing common resilience mechanisms to both drought and waterlogging.

The energy shortage caused by waterlogging also leads to significant changes in sugar metabolism. The accumulation of soluble sugars such as glucose and fructose, might be a consequence of sink-source imbalances during waterlogging and thereby, a decline in shoot-to-root sugar transport. Strikingly, we observed upregulation of *SP6A,* a positive regulator of tuberisation, thus suggesting potential roles of this gene in short-term responses to waterlogging.

Prolonged exposure to waterlogging revealed several aspects of late responses and factors contributing to potato susceptibility to waterlogging. Leaves of waterlogged plants overcome energy shortages by recycling carbon from amino acids and GABA. The latter, plays an important role not only in TCA replenishment but also in ion homeostasis and reduction of oxidative stress (Lothier et al., 2020; Wu et al., 2021). We observed a strong increase in free amino acids that, together with the upregulation of POP2 and an aminopeptidase (Figure 6), suggests increased protein breakdown and utilization of amino acids as alternative energy sources. Furthermore, the downregulation of RPL27 (and other ribosomal proteins) could indicate the shutting down of energy-demanding processes, such as protein synthesis, as a response to this energy shortage.

Potato susceptibility to prolonged waterlogging was evidenced by other multi-level events such as the upregulation of proteins related to protein and cell wall component turnover, *RboHA* upregulation and photosynthesis impairment (Figure 2, Supp. Figure 3, Figure 6). It is also explained by increased ABA signalling and biosynthesis and *RBohA* expression, which convergently indicate increased tissue dehydration and oxidative stress that is reflected in the HTP data (Figure1C, Figure 2). This includes decreased tuber number and weight, indicating a retardation of both tuber initiation and bulking. As tuberisation is a particularly energetically expensive process, the imposition of root zone hypoxia likely disrupts the underground sink force essential for stolon development, tuber initiation and bulking.

Altogether, our data suggest that two weeks of waterlogging led to near-lethal effects, and even if acclimation responses were activated, overall, they could not compensate for maintaining root function (i.e. unrecovered water consumption, Supp. Table 4) and general plant survival, even during recovery, thus confirming the high susceptibility of potato to waterlogging.

### Conclusion

The present comprehensive approach produced a rich integrated dataset, which enabled diverse exploration of molecular mechanisms across various levels and processes. Through the connection of phenotype to molecular responses, we attained deeper insights into the intricate regulation of metabolic and phenotypic traits. This should now guide the identification of key regulators that govern the interplay between molecular dynamics and their phenotypic expressions. The utilization of both knowledge-based approaches and multivariate statistical methods played a crucial role in deciphering complex molecular regulatory networks and their association with phenotypic and physiological traits, thereby facilitating the rapid generation of hypotheses.

In addition to several novel insights into potato stress responses, this study also provides a blueprint for performing and analysing single and multiple stress and effective integration of large datasets for potato. Importantly, this setup could be applied also for other plant species. These advancements hold significant implications for potato breeding strategies, providing a deeper understanding of plant stress responses and expediting trait selection. As agricultural landscapes confront challenges like climate change and population growth, embracing multi-omics integration holds promise for cultivating resilient potato varieties that can thrive in various conditions.

## METHODS

### Plant growth conditions and sampling

150 in-vitro potato cuttings (*Solanum tuberosum* cv Desirée) were cultivated and grown as described in the supplementary methods. After 32 days of cultivation, plants were randomly distributed into 6 groups (6 plants each) referring to control group and 5 different stress conditions (heat, drought, combined heat and drought, waterlogging and combination of heat, drought and waterlogging) (Figure 1). Plants were moved into two growth units of Growth Capsule (PSI; (Photon Systems Instruments), Czech Republic) where climate conditions for day/night temperature were set in one unit to 22/19°C, referring to control conditions, and in the second unit to 30/28 °C, referring to heat conditions. In both units growing light intensity was set at 330 µmol m^-2^ s^-1^ PPFD and relative humidity was maintained at 55%. All plants were measured under control in day 0 then the stress treatments depicted schematically in Figure 1. The treatments were applied as the following: (1) Control conditions – cultivation at 22/19 °C, watering up to 60% FC; (2) Drought conditions – cultivation at 22/19 °C, watering up to 60% FC until day 7, then reduce watering to 30% FC for 1 week (until day 14); (3) Heat conditions – cultivation at 30/28 °C for 2 weeks, watering up to 60% FC until day 14; (4) Heat + Drought conditions - cultivation at 30/28 °C for 2 weeks, watering up to 60% FC for 1 week (until day 7), then reduce watering to 30% FC for 1 week (until day 14); (5) Waterlogging conditions – cultivation at 22/19 °C, watering up to 130% FC for 2 weeks (until day 14); (6) Heat + Drought + Waterlogging conditions – cultivation at 30/28 °C for 2 weeks with watering up to 60% FC for 1 week (until day 7), then reduce watering to 30% FC until day 14 followed by inducing waterlogging by cultivation at 22/19 °C for 1 week with watering up to 130% FC until day 21. Except for Heat + Drought + Waterlogging conditions, all stress treatments were followed by one week of recovery (from day 15 and until day 21) in control conditions.

Plants were divided into two sets, “phenotyping plants” and “plants for tissue harvest” (see Supp. Table 1). Phenotyping set consisted of 6 replicates per treatment, in total 36 plants, and was used for daily image-based phenotyping (for definition of scored traits see Supp. Table 2).

### High-throughput phenotyping

Prior to the stress treatment initiation and during the stress treatments, all plants were daily phenotyped. A comprehensive phenotypical protocol was used for the acquisition of physiological and morphological traits according to the described method (Abdelhakim et al., 2024). All imaging sensors for digital analysis are being implemented in the PlantScreen^TM^ Modular system (PSI, Czech Republic). The photosynthesis□related traits were determined using kinetic chlorophyll fluorescence imaging where the selected protocol for measuring plants was similar to the defined approach (Abdelhakim et al., 2024). The measurement of temperature profiles of the plants was measured using thermal imaging, where the acquisition and segmentation of the images were processed as described in (Abdelhakim et al., 2021; Findurová et al., 2023). The morphological and growth dynamics were determined using both top and multiple angles (0°, 120°, and 240°) side view RGB imaging, and images were processed as described by (Awlia et al., 2016). Each pot was loaded onto a transport disk automatically moving on a conveyor belt between the automatic laser height measuring unit, acclimation unit, robotic-assisted imaging units, weighing and watering unit, and the cultivation greenhouse located area. The raw images were automatically processed and parameters were extracted through PlantScreen^TM^ Analyzer software (PSI, Czech Republic) (Supp. Table 2). Statistical evaluation was performed to check the differences between the treatments using Wilcoxon test (Supp. Table 3).

### Tissue sampling

Leaf sampling was conducted on days 1, 7, 8, 14, 15, and 21 after stress treatment initiation (Treatment days). The 2^nd^ and 3^rd^ fully-developed leaves were harvested and flash-frozen in liquid nitrogen. Subsequently, leaf tissue was homogenised, aliquoted and distributed for individual follow-up transcriptomics, metabolomics, hormonomics, and proteomics analyses. Remaining above ground tissue was harvested and total fresh weight (FW) and dry weight (DW) was defined (Supp. Table 4). In total 112 plants with 4 replicates per sampling time and per treatment were collected. The 4^th^ leaf was harvested as well to calculate relative water content (RWC) (Supp. Table4), and three leaf disks were collected and weighed, then soaked in water to determine the turgor weight (TW) and dried in the oven to calculate RWC (Supp. Table 2). In addition, at the end of the experiment, the below ground tissue was collected, where number of tubers per plant and total weight were assessed from four replicates per treatment. Harvest index was calculated as a ratio between tuber weight and the total biomass.

### Multi-omics analysis

#### Transcriptomic marker analysis

RT-qPCR was performed to assess the expression of 14 marker genes involved in redox homeostasis) hormonal signalling - ethylene, abscisic acid, cytokinin, salicylic acid, jasmonic acid, heat stress, tuber development, circadian clock, and calcium signalling using previously validated reference genes (Supp. Table 7).

RNA was extracted and DNase treated using Direct-zol RNA Miniprep Kit (Zymo Research, USA) from 80-100 mg of frozen homogenised leaf tissue, followed by reverse transcription using High-Capacity cDNA Reverse Transcription Kit (Thermo Fisher, USA). The expression of the target and reference genes was analysed by qPCR, as described previously (Petek et al., 2014; Abdelhakim et al., 2021). QuantGenius (http://quantgenius.nib.si), was used for quality control, standard curve-based relative gene expression quantification and imputation of values below level of detection or quantification (LOD, LOQ) (Baebler et al., 2017).

#### Hormonomics

Concentration of the endogenous abscisate metabolites, auxin metabolites, jasmonates and salicylic acid were determined in 10 mg of frozen homogenised leaf tissue according to the method described (Flokova et al., 2014) and modified by Široká et al. (Siroka et al., 2022) (see Supplementary methods for details). All experiments were repeated as four biological replicates.

#### Metabolomics

For determination of soluble sugar, starch and amino acid contents, 30 - 50 mg of freeze-dried leaf or tuber material were extracted with 1 ml of 80% (v/v) ethanol. Soluble sugar and starch content was determined as described in Hastilestari et al. (2018), while amino acids sample preparation and measurements were performed as described elsewhere (Obata et a. 2020). (Smith and Zeeman, 2006) (Obata et al., 2020).

#### Proteomics

High-throughput shotgun proteomics was done according to (Hoehenwarter et al., 2008) with following modifications: 40 mg of leaf tissue from multiple stress conditions were freeze-dried in liquid N_2_ and ground using mortar and pestle. The proteins were extracted, pre-fractionated (40µg of total protein were loaded onto the gel (1D SDS-PAGE), trypsin digested and desalted (using a C18 spec plate) according to a previously described method (Chaturvedi et al., 2013; Ghatak et al., 2016). One µg of purified peptides was loaded onto a C18 reverse-phase analytical column (Thermo Scientific, EASY-Spray 50 cm, 2 µm particle size). Separation was achieved using a two-and-a-half-hour gradient method, starting with a 4–35% buffer B (v/v) gradient [79.9% ACN, 0.1% formic acid (FA), 20% ultra-high purity water (MilliQ) over 90 minutes. Buffer A (v/v) consisted of 0.1% FA in high-purity water (MilliQ). The flow rate was set to 300 nL/min. Mass spectra were acquired in positive ion mode using a top-20 data-dependent acquisition (DDA) method. A full MS scan was performed at 70,000 resolution (m/z 200) with a scan range of 380–1800 m/z, followed by an MS/MS scan at 17,500 resolution (m/z 200). For MS/MS fragmentation, higher energy collisional dissociation (HCD) was used with a normalized collision energy (NCE) of 27%. Dynamic exclusion was set to 20 seconds.

Raw data were searched with the SEQUEST algorithm present in Proteome Discoverer version 1.3 (Thermo Scientific, Germany) described previously (Chaturvedi et al., 2015; Ghatak et al., 2020). Pan-transcriptome (Petek et al., 2020) protein fasta was employed. The identified proteins were quantitated based on total ion count and normalised using the normalised spectral abundance factor (NSAF) strategy (Paoletti et al., 2006).

#### Data analysis

The programming environments R v.4.3 and v4.4 (https://www.r-project.org/) and Python v3.8 (www.python.org) were used. Experimentally acquired data and data required to reproduce the analysis are available from Supplemental Table 4 and NIB’s GitHub repository (https://github.com/NIB-SI/multiOmics-integration) were used.

#### Data preprocessing

A master sample description metadata file was constructed (Supp. Table 1). Potential inconsistencies between replicates were examined using pairwise plots between omics levels, multidimensional scaling plots and scatterplot matrices within omics’ levels using the vegan v2.6.-4 R package (Oksanen et al., 2022). Missing values were handled as described in the Supplementary methods. Due to many missing values, the neoPA (hormonomics) variable was excluded from further analysis.

Variable selection was conducted on the non-invasive phenomics variable sets (Supp. Table 4). The random forest (RF) algorithm from the R package caret v6.0-94 (Kuhn, 2008) as well as the python package scikit-learn v1.2.0 were used with default settings, as RF showed the best performance out of a selection of algorithms. Recursive feature elimination was applied in R and multiple importance scores, including mutual information, Anova, RF importance and SHAP values (Lundberg and Lee, 2017) were computed in Python, showing consistencies between the approaches for the top 5 variables (Top area, compactness, qL_Lss, ΔT, water consumption; nonredundancy ranking in R). The sixth variable (Fv/Fm_Lss) was selected based on expert knowledge.

Gene set enrichment was performed on the proteomics dataset using GSEA v4.3.2 (Subramanian et al., 2005) and in-house generated gene sets (Supp. File 3, Supp. Table 6) and visualised using biokit v0.1.1. Proteomics differential expression was conducted using the DEP v 1.22.0 package (Zhang et al., 2018) (Supp. Table 5). For downstream proteomics analyses, differentially abundant and enriched proteins (from pathways important for this experimental setup) were used. Waterlogging stress was cut-off at one-week duration, while triple stress (HDW) was not considered in downstream analyses due to poor plant performance.

#### Analysis of individual omics data layers

Pearson correlation coefficient (PCC) heatmaps (pheatmap v1.0.12, heatmaply v. 1.5.0) were generated within each treatment and for explicit treatment duration. Permutation-based t-test (MKinfer v1.2) was used to denote differences between specific treatment and control within the corresponding time-point (Kohl, 2024). Corresponding log2FC were calculated. For downstream analyses, 4 out of 6 replicates were chosen from non-invasive phenomics measurements to allow integration with invasive phenomics and other omics measurements conducted on 4 replicates.

#### Integration across different omics datasets

Correlations between components measured in various Omics’ levels were calculated and visualised using DIABLO (Singh et al., 2019) as implemented in the mixOmics v6.24.0 package (Rohart et al., 2017). The correlation matrix was calculated separately for each stress as well as for control.

#### Integration of data with prior knowledge

A background knowledge network was manually constructed considering biochemical pathways between measured variables. Where necessary, pathways were simplified to only include representative variables, to prevent addition of many unmeasured nodes that would impede the visualisation. Proteomics differential expression results were merged with t-test and log2FC results (Supp. Table 3). Final networks were visualised using DiNAR (Zagorscak et al., 2018) and Cytoscape (Shannon et al., 2003).

For additional reports and some results not used in this manuscript see supplementary methods and a project’s GitHub repository https://github.com/NIB-SI/multiOmics-integration.

## Acknowledgements

The authors wish to thank Marijke Woudsma and Doretta Boomsma from HZPC Research for providing the Desiree plantlets for this study and Mirella Sorrentino for help with conducting the experiments at Photon Systems Instruments (PSI) Research Center (Drásov, Czech Republic). Moreover, we the thank following colleagues for their assistance and expertise: S. Reid (FAU) and D. Pscheidt (FAU) for measuring metabolite content, and Katja Stare (NIB) and Nastja Marondini (NIB) for measuring gene expression.

## Author contributions

KP, KG, SS, RS, CBa and MT conceptualized the study and designed the experiments, KP measured phenotypic traits, ŠB measured gene expression, ON, AP and JŠ measured hormone content, CS measured metabolite content, PC, AG prepared samples and performed proteomics analysis, LAS performed LC-MS measurements for proteomics analysis; MZ curated data; MZ, CB, AB, AZ and KG defined formal analysis methodology; MZ, CB, JZ, and AB conducted statistical, mathematical and computational analysis; MZ, LA, NR, CB, AB, ŠB and KG performed data visualisation; MZ, LA, NR, CB, SS, RS, KG, CBa and MT wrote the original draft. All authors edited the manuscript and approved the final manuscript version. Detailed author contributions are available from the Supp. Table 8.

## Supplementary data

**Supplementary File 1: Principal coordinates analysis (PCoA) of Bray-Curtis dissimilarity between samples.** Interactive 3D plot shows mapping to three dimensions. Spheres are coloured by condition. The numerical values used to construct this figure can be found in Supplementary Table 4. Variables were min-max scaled.

**Supplementary File 2: Differential Network Analysis animation.** Export from DiNAR app (Zagorščak et al., 2018). Node colours correspond to upregulation (red) and downregulation (blue) compared to control (p-value adjusted < 0.05). The size of nodes corresponds to absolute log2FC values. Order of visualisation: 0: background network, 1: 1 day of drought, 2: one week of drought 3: one day of heat, 4: one week of heat, 5: 8 days of heat, 6: two weeks of heat, 7: 8 days of heat + one day of drought, 8: two weeks of heat + one week of drought, 9: one day of waterlogging, 10: one week of waterlogging.

**Supplementary File 3: Pan-transcriptome Gene Matrix Transposed (GMT) file format.** Tab delimited file utilised for gene set enrichment analysis that describes gene sets from pan-transcriptome of three potato genotypes: Desiree, PW363 and Rywal (Petek et al., 2020).

**Supplementary Figure 1: Plant morphological responses of control and stress-treated plants**. (A) Top view RGB images at selected time points of tissue sampling starting from day 1 where i) in the first week heat stress was induced for H, HD, HDW marked rows, then ii) in the second week drought was induced for D, HD, HDW up to day 14, and finally iii) in the third week waterlogging was induced for HDW up to day 21, while the other treatments were recovered. (B-E) Parameters based on analysed RGB images; Top area, plant height, compactness, and relative growth rate (RGR). The data represent mean values ± standard error of mean (S.E.M) (n = 6).

**Supplementary Figure 2: Log2FC of a tuber metabolite’s relative abundance between stress and control.** Numbers in bold correspond to comparisons with p-value < 0.05. Additionally, log2FC for number of tubers and total tuber weight is shown.

**Supplementary Figure 3: Correlation analysis within and between omics levels.** For correlation analysis within omics levels heatmaps display Pearson correlation coefficient (PCC). For analysis between components of different molecular levels heatmaps display canonical correlation analysis (CCA) results. Variable prioritisation was conducted using multiblock sPLS-DA (Singh et al., 2019).

**Supplementary Table 1: Phenodata.** Experimental design and days of tissue sampling.

**Supplementary Table 2: Phenomics featuredata.** Phenotyping trait description from multiple imaging sensors and traits from the final destructive harvest.

**Supplementary Table 3: Statistical results across different omics levels.** Combined table contains statistical information and various annotations for molecules shown in biochemical knowledge network (see Figure 2,4,5,6 and Supplementary File 2).

**Supplementary Table 4: Omics measurements**. Multi-sheet file containing tuber (metabolomics, harvest information) and leaf data (phenomics, transcriptomics, hormonomics and metabolomics). Phenomics data is separated to non-invasive and invasive trait measurements. For trait description see Supplementary Table 2.

**Supplementary Table 5: Differential protein expression.** Results table of differential protein expression from R package DEP (Zhang et al., 2018) complemented by MapMan annotations (Ramšak et al., 2014) and Arabidopsis thaliana orthologue identifiers, short names, and gene descriptions (Bleker at al., 2024, Zagorščak et al., 2018).

**Supplementary Table 6: Gene set enrichment analysis of proteomics data.** MapMan v3 pathway enrichment results containing classes of proteins that are over-represented in stress treatments compared to control (FDR cut-off 0.05 and 0.1). MapMan pathways are defined in supplementary file 3.

**Supplementary Table 7: Genes used for transcriptome analysis using quantitative PCR.** Gene functional group, name (abbreviation) and description, as well as corresponding primer and probe sequences, source and amplification efficiency are shown. Suitability of reference genes was validated using RefFinder (Xie et al., 2023).

**Supplementary Table 8: CRediT (Contributor Roles Taxonomy) authorship statement.**

## Competing interests

The authors declare no competing interests.

## Funding

This work was funded by the EU H2020-SFS-2019-2 RIA project ADAPT, GA 2020 862-858 and the Slovenian Research Agency (ARIS) under grant agreements P4-0165, J2-3060, and Z4-50146. Moreover, this work was partially supported by the Ministry of Education, Youth and Sports of the Czech Republic with the European Regional Development Fund-Project “SINGING PLANT” (no. CZ.02.1.01/0.0/0.0/16_026/0008446).

## Data availability statement

Experimentally acquired data and data required to reproduce the analysis are available from Supp. Table 4 and NIBs’ GitHub repository https://github.com/NIB-SI/multiOmics-integration.

The MS/MS spectra of the identified proteins and their meta-information from both databases have been deposited to the ProteomeXchange Consortium via the PRIDE partner repository (https://www.ebi.ac.uk/pride) with the dataset identifier PXD052587.

## Notes

### Competing Interest Statement

The authors have declared no competing interest.

### Summary of Updates

The manuscript was carefully streamlined and shortened. Parts of the Materials and Methods section into a supplementary text. Statistical analysis and machine learning details are now part of the methods section and supplementary methods section. DIABLO analysis details and downstream procedures can be found in supplementary methods section. We have completely re‐written the abstract and introduction and have also re‐structured the entire manuscript text to better connect the findings and to focus on the main parts. Statistical procedures and detail p‐values that support the claims about significance in the results section are explained as well as the reason for choice of random forest as a model to assess variable importance and its effects on the downstream findings. We discussed the fact that upregulation of enzymes and changes in metabolite levels are supported by correlations analysis and proteomics as well as phenomics data to underpin the added value of the multi‐level analysis in all respective discussions. We further discussed the increased JA levels and related proteins in more detail. Therefore we conclude with emphasizing the need to evaluate potato varieties tolerant to thermal stress to discover if JA metabolism would play a significant role in thermo‐tolerance. We detailed specifically the suitability of the developed multi‐omics pipeline compared to other existing tools. We did extensive analysis between different molecular levels using DIABLO. The procedure is described in the Supplementary Methods. We clearly specified the accessibility of the data in the public repositories and opened the proteomics data for open access. High resolution images in vector formats and all supplemental data to the project are now available in the GitHub repository. https://github.com/NIB-SI/multiOmics-integration/.

