## Supplemental Figures 1-3 for "Integration of multi-omics data and deep phenotyping of potato enables novel insights into single- and combined abiotic stress responses"

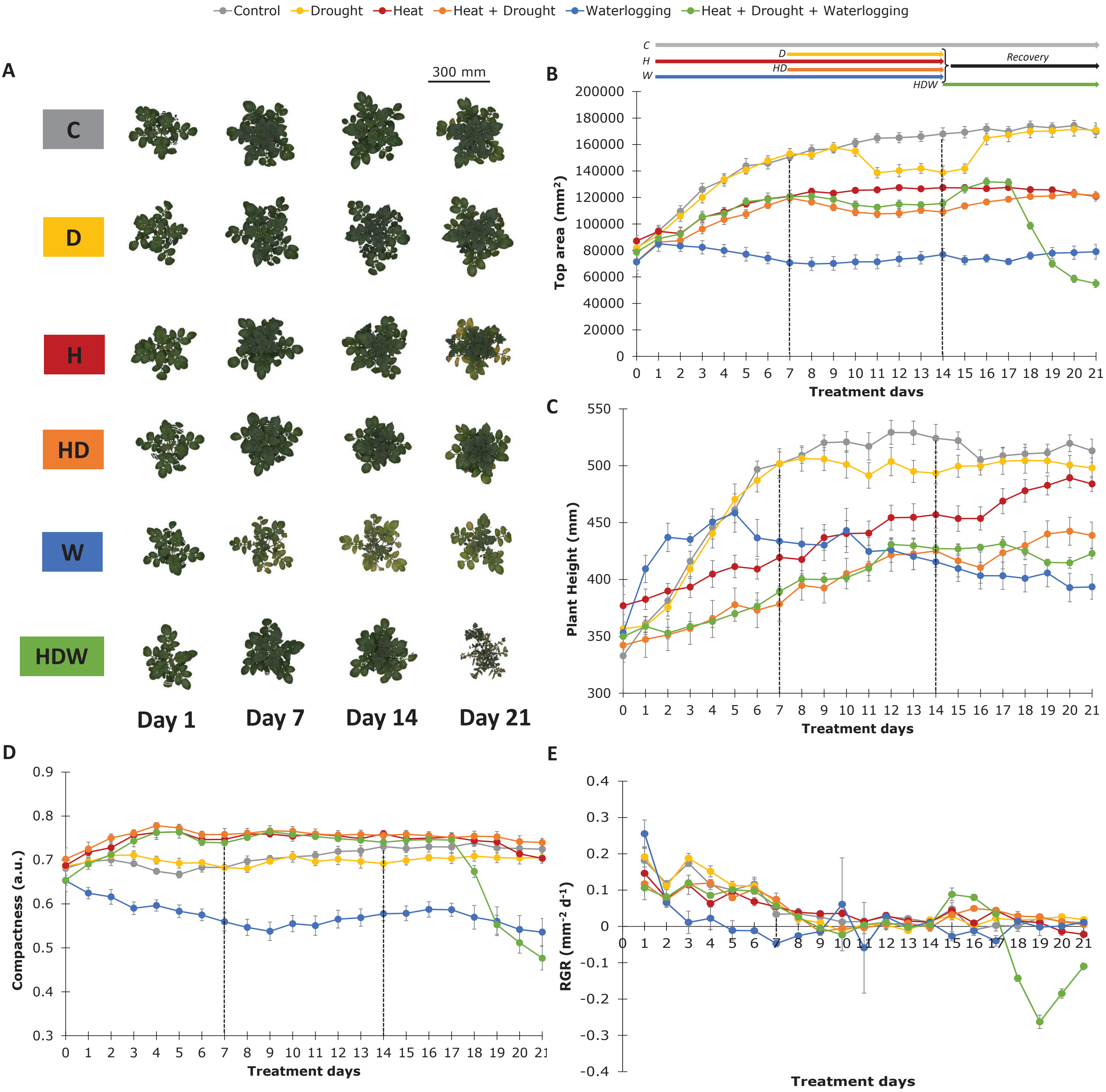


**Supplementary Figure 1: Plant morphological responses of control and stress-treated plants.** (A) Top view RGB images at selected time points of tissue sampling starting from day 1 where i) in the first week heat stress was induced for H, HD, HDW marked rows, then ii) in the second week drought was induced for D, HD, HDW up to day 14, and finally iii) in the third week waterlogging was induced for HDW up to day 21, while the other treatments were recovered. (B-E) Parameters based on analysed RGB images; Top area, plant height, compactness, and relative growth rate (RGR). The data represent mean values ± standard error of mean (S.E.M) (n = 6).

| | D28 | H28 | HD28 | W28 |
|---|---|---|---|---|
| Glucose | -0.520 | **0.886** | **1.026** | -0.363 |
| Fructose | 0.185 | -0.362 | 0.497 | 0.708 |
| Sucrose | 0.014 | 0.019 | 0.480 | **2.050** |
| Starch | 0.115 | **-0.419** | **-0.289** | **-0.563** |
| Ala | -0.137 | **-0.731** | -0.437 | 0.919 |
| Arg | -0.291 | **-0.710** | -0.262 | **1.588** |
| Asn | -0.124 | -0.508 | -0.187 | **2.802** |
| Asp | -0.107 | -0.038 | 0.164 | **1.230** |
| Gln | -0.056 | -0.562 | -0.098 | **2.611** |
| Glu | -0.225 | **-0.404** | **-0.299** | 0.091 |
| Gly | -0.300 | **-0.846** | -0.568 | **1.045** |
| His | **-0.428** | **-0.664** | -0.322 | **1.546** |
| Ile | -0.250 | -0.542 | -0.323 | **1.001** |
| Leu | -0.298 | -0.295 | -0.075 | **1.017** |
| Lys | **-0.477** | **-0.876** | **-0.753** | 0.434 |
| Met | -0.239 | **-0.817** | **-0.485** | **0.408** |
| Phe | -0.389 | **-1.437** | **-1.090** | **-1.158** |
| Pro | -0.241 | -0.131 | 0.028 | **2.850** |
| Ser | -0.146 | -0.595 | -0.374 | **1.447** |
| Thr | -0.083 | **-0.775** | -0.365 | **1.189** |
| Tyr | -0.361 | **-0.824** | **-0.742** | -0.059 |
| Val | -0.141 | **-0.640** | -0.301 | **0.963** |

| | D28 | H28 | HD28 | W28 |
|---|---|---|---|---|
| Number of tubers | +26% | **+32%** | +16% | **-82%** |
| Total tubers weight | -4.6% | **-48%** | **-48%** | **-97%** |

**Supplementary Figure 2: Log2FC of a tuber metabolite's relative abundance between stress and control.** Numbers in bold correspond to comparisons with p-value < 0.05. Additionally, log2FC for number of tubers and total tuber weight is shown.

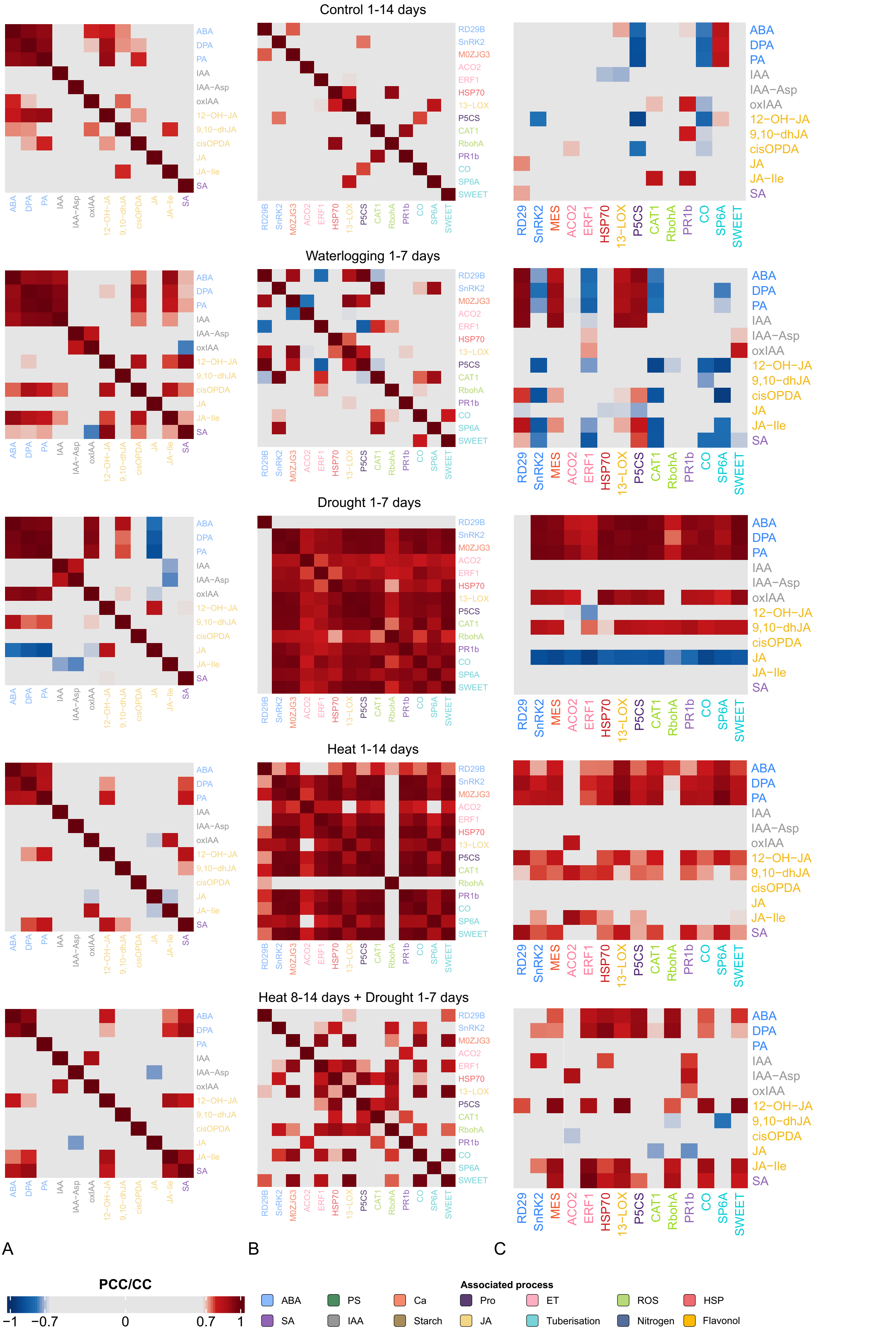


**Supplementary Figure 3: Correlation analysis within and between omics levels.** For correlation analysis within omics levels heatmaps display Pearson correlation coefficient (PCC). For analysis between components of different molecular levels heatmaps display canonical correlation analysis (CCA) results. Variable prioritisation was conducted using multiblock sPLS-DA (Singh et al., 2019).
